# DNA Damage and Repair Mechanisms in Duckweed (*Spirodela polyrhiza*) Under Ultraviolet-B (UV-B) Light Stress

**DOI:** 10.64898/2026.09.02.748994

**Authors:** Manvitha A. Sanjaya, Shivasharanappa S. Patil

## Abstract

Exposure to Ultraviolet-B (UV-B) light can adversely affect plant growth and cellular integrity by inducing oxidative stress and DNA damage. In this study, we investigated UV-B-induced DNA damage and repair responses in the aquatic monocotyledonous plant species *Spirodela polyrhiza* (duckweed). We exposed 13-day-old duckweed plantlets to broadband UV-B light for 1-10 min, followed by recovery periods of up to 24 h under normal growth conditions. We observed progressive chlorosis, wilting, and diminished plant vigor with longer duration of UV-B light exposure. Agarose gel electrophoresis demonstrated compromised genomic DNA integrity immediately after UV-B light treatment, with partial restoration of DNA quality during recovery. Immuno-slot blot assays established the accumulation of two major UV light–induced photoproducts, cyclobutane pyrimidine dimers (CPDs) and 6-4 pyrimidine-pyrimidone photoproducts [(6-4)PPs] in a dose-dependent manner following UV-B light exposure. Notably, the abundance of these DNA lesions declined substantially after recovery, indicating activation of endogenous DNA repair mechanisms. Staining with 3,3′-diaminobenzidine revealed elevated accumulation of hydrogen peroxide immediately following UV-B exposure, suggesting enhanced oxidative stress. Collectively, these findings demonstrate that *S. polyrhiza* possesses efficient mechanisms for sensing, repairing, and mitigating DNA damage induced by the oxidative stress resulting from UV-B light exposure. This study highlights the potential of duckweed as an effective model system for investigating DNA damage and repair pathways under UV-B light stress in plants.

## Introduction

Sunlight is vital for life on Earth. For instance, humans require sunlight to produce vitamin D. Photosynthetic organisms, such as plants and microalgae, depend on sunlight to produce energy to support cellular metabolism through photosynthesis. In addition to serving as a primary source of energy, sunlight is also a significant source of ultraviolet (UV) light. UV light radiation is characterized by its wavelengths and properties, namely UV-A (320-420 nm), UV-B (280-320 nm), and UV-C (180-280 nm) [1]. UV-A and UV-B light penetrate the ozone layer and reach the Earth’s surface, whereas the Earth’s atmosphere reflects almost all UV-C light [2,3].

UV-B light, a component of sunlight, can have both beneficial and harmful effects on living organisms. In humans, exposure to UV-B light primarily affects the epidermis, the outermost layer of the skin. Excessive exposure to UV-B light can damage cellular DNA, proteins, and lipids by generating reactive oxygen species (ROS) [4]. This damage can lead to sunburn, premature skin aging, inflammation, immune suppression, and an increased risk of skin cancers, including melanoma and non-melanoma skin cancers [3]. At the same time, exposure to UV-B light also stimulates melanin production as a protective response [5,6].

In plants, UV-B light mainly affects epidermal and mesophyll cells. Plant epidermal tissues act as the first protective barrier against UV-B damage by accumulating UV-absorbing compounds, such as flavonoids and phenolics [7]. Excessive exposure to UV-B light can impair photosynthesis, damage DNA, disrupt membrane integrity, lower chlorophyll content, and inhibit plant growth and productivity [8]. At the cellular level, UV-B light induces oxidative stress through ROS production, leading to the damage of nucleic acids, proteins, and cellular membranes [9]. However, plants also possess adaptive and protective mechanisms, including activation of antioxidant enzymes, biosynthesis of protective pigments, and UV RESISTANCE LOCUS 8 (UVR8-mediated signaling, antioxidant responses, and DNA-repair pathways that together promote acclimation to UV-B stress [10,11,12].

In both humans and plants, the epidermis is rich in chromophores, a type of molecule that can absorb light energy including UV-B light [13,14,15]. DNA itself functions as a chromophore, as its constituent pyrimidine bases strongly absorb UV light [13], disrupting the double bonds within pyrimidine bases, particularly cytosine and thymine, leading to the formation of DNA lesions known as pyrimidine dimers [11,16]. The two major UV light-induced photoproducts are cyclobutane pyrimidine dimers (CPDs) and 6-4 pyrimidine-pyrimidone photoproducts [(6-4)PPs] [17,18]. In mammals, these lesions are primarily repaired through the nucleotide excision repair (NER) pathway [19,20,21].

Because sunlight is indispensable for photosynthesis, plants cannot avoid UV light exposure and have therefore developed efficient DNA repair systems to maintain genome stability [22]. In addition to NER, plants use photoreactivation, base excision repair (BER), translesion synthesis, and double-strand-break repair pathways to maintain genome stability after UV stress [23,24,22].

Notably, plant genomes harbor homologs for DNA repair genes previously characterized in yeast (*Saccharomyces cerevisiae*) and mammals [12,23]. Consequently, plants have emerged as valuable model systems for investigating genes and pathways involved in DNA damage sensing and repair. Previous studies have identified numerous genes associated with DNA repair mechanisms in model plant species such as Arabidopsis (*Arabidopsis thaliana*) [25,26,27]. Recent work shows that plant DNA-damage responses are tightly coupled to cell-cycle checkpoints, transcriptional reprogramming, and repair-pathway activation serving as important regulatory nodes [23,24]. However, limited information is available regarding the conservation and function of these pathways in other plant species.

In this study, we investigated UV-B light-induced DNA damage and repair mechanisms in the duckweed *Spirodela polyrhiza. S. polyrhiza* is a rapidly reproducing aquatic monocot with a small genome and increasingly well-resolved chromosome-scale genomic resources, making it an attractive model for molecular and physiological studies [28,29,30]. These characteristics provide an excellent platform for studying UV-B light-induced DNA damage and repair mechanisms as an alternative to mammalian model systems.

## Materials and Methods

### Maintenance of Spirodela polyrhiza cultures

Three to four frond colonies of an *S. polyrhiza* wild-type strain were inoculated into Petri dishes containing semi-solid half-strength liquid Schenk and Hildebrandt (SH) basal salt mixture (0.8 g/L; Sigma Cat. No. S6765; PhytoTechnology Laboratories Cat. No. S816) supplemented with 0.5% (w/v) sucrose. Cultures were maintained for 13 days in a controlled plant growth chamber under a 16-h light/8-h dark photoperiod with a light intensity of 80 µmol m^−2^ s^−1^ at 25°C for subsequent experiments.

### UV-B light treatment and recovery

UV-B treatments were performed using a QUV Gen 4 SOLAR EYE chamber (Model QUVSE120G4; Q-Lab Corporation, Westlake, OH, USA) equipped with UVB-313 EL lamps. The UV-B irradiance was calibrated using a Legacy CR10 UVA/UVB ultraviolet radiometer (QLab Corporation) according to the manufacturer’s specifications and adjusted to 0.2 μmol photons m^−2^ s^−1^ at the sample exposure plane. The QUV chamber was maintained at 25°C during irradiation.

Approximately 13–day-old *Spirodela polyrhiza* cultures maintained on SH medium were used for UV-B exposure experiments. Cultures were grown in Petri dishes at 25°C under a 16-h light/8-h dark photoperiod in a controlled-environment plant tissue-culture chamber (Percival Scientific, USA). To minimize potential differences in temperature associated with the transition from the growth chamber to the UV-B exposure system, Petri-dish lids were removed approximately 24 h before irradiation, and the cultures were maintained under the same environmental conditions until treatment.

For UV-B exposure, individual Petri dishes containing duckweed plantlets were positioned horizontally within adjustable quadrant boxes (Q-Lab Corporation). The open side of each quadrant box was oriented toward the UV-B EL lamps, and the sample surface was positioned approximately 12 inches (30.5 cm) from the UVB-313 EL lamps. The Petri dishes remained open during irradiation to avoid attenuation or spectral modification of the incident UV-B radiation by the lids. Duckweed cultures were exposed to UV-B light for 1, 2.5, 5, or 10 min, corresponding to increasing UV-B exposure durations at the calibrated irradiance. Following irradiation, the plantlets were immediately returned to the controlled-environment plant tissueculture chamber and allowed to recover under the standard growth conditions described above (25°C, 16-h light/8-h dark photoperiod).

For each exposure duration, UV-B-unexposed cultures maintained under identical growth conditions served as untreated controls. Control Petri dishes were handled in parallel with the treated samples but were not exposed to UV-B radiation. Plant material was collected at 0, 12, and 24 h after UV-B exposure for subsequent analyses of UV-B-induced DNA damage, DNA repair-associated responses, oxidative stress, and gene expression. The 0-h samples were collected immediately following completion of the respective UV-B treatment and before the recovery period, whereas the 12- and 24 h samples represented recovery time points under the standard growth conditions.

### Genomic DNA isolation and gel electrophoresis

For genomic DNA extraction, samples were collected from duckweed cultures immediately after exposure to UV-B light and after 12 h or 24 h of recovery, immediately frozen in liquid nitrogen, and stored at −80°C until analysis. Genomic DNA was isolated from three biological replicates using a DNeasy Plant Mini Kit (Qiagen, USA; Cat. No. 68163) according to the manufacturer’s instructions with minor modifications.

Briefly, up to 0.1 g fresh weight or 0.02 g lyophilized tissue per sample was homogenized using a TissueRuptor, TissueLyser, or mortar and pestle. Then, samples were transferred to centrifuge tubes containing 400 mL of preheated Buffer AP1 and 4 μL RNase A, followed by vigorous vortexing until complete tissue disruption was achieved. Samples were incubated at 65°C for 10 min with inversion every 2-3 min. Subsequently, 130 μL of Buffer P3 was added to each sample, and the mixtures were incubated for 5 min on ice to facilitate precipitation of proteins and secondary metabolites. Following centrifugation at 3,000-5,000 *g* for 5 min at room temperature (RT), supernatants were transferred onto QIAshredder spin columns placed in 50-mL collection tubes and centrifuged at 20,000 *g* for 2 min.

The clarified lysate was transferred into fresh tubes and mixed immediately with 1.5 volumes of Buffer AW1. These mixtures (maximum 650 μL) were transferred into DNeasy Mini spin columns placed in 2-mL collection tubes. The tubes were centrifuged at 6,000 *g* for 1 min, and the flow-through was discarded. This step was repeated. The columns were then washed with 500 μL Buffer AW2 and centrifuged for 2 min at 20,000 *g*. The flow-through was discarded. For DNA elution, the spin columns were transferred to new 2-mL tubes and incubated with 100 μL Buffer AE for 5 min at room temperature (25°C), prior to centrifugation at 6,000 g for 1 min. The DNA elution step was repeated to maximize DNA recovery. DNA concentration was determined using a PicoGreen dsDNA assay and the genomic DNA was stored at −20°C until use. DNA integrity was evaluated by gel electrophoresis of 2 µL genomic DNA on a 1% (w/v) agarose gel run at 100 V for about 30 min. Following electrophoresis, gels were stained with SYBR Safe DNA Gel Stain (Thermo Scientific, USA; Cat. No. S33102), and images were acquired using a Bio-Rad ChemiDoc Imaging System.

### Slot-blot analysis of UV light–induced DNA lesions

Slot-blot analysis was performed to quantify UV light-induced DNA lesions, including CPDs, (6-4)PPs, and platinum-induced adducts [Pt-(G-G/C-G)]. DNA concentrations were determined using a PicoGreen dsDNA assay, and equal amounts of DNA were diluted in 300 µL nucleasefree water. Typically, 150 ng DNA was used for CPD detection, whereas 300-500 ng DNA was used for detection of Pt-(G-G/C-G) adducts and (6-4)PPs. DNA samples were denatured at 100°C for 10 min and immediately cooled on ice for 10 min. Equal volumes of ice-cold 2 M ammonium acetate (pH 7.0) were then added to neutralize the samples.

A slot-blot manifold apparatus (96-well Bio-Dot; Bio-Rad, USA; Cat. No. 1706545) was assembled and operated according to the manufacturer’s instructions. Three sheets of blotting paper and a 0.45-µm nitrocellulose membrane were equilibrated in 6× SSC buffer for 10 min before assembly. Samples were loaded onto the membrane under gentle vacuum pressure according to a predetermined blot map [31]. Following sample application, membranes were washed with Tris-EDTA (TE) buffer and subsequently with 2× SSC buffer. The washed membranes were baked in a vacuum oven at 80°C for 2 h and then blocked in phosphate-buffered saline (PBS) containing 5% (w/v) non-fat dry milk and 0.1% (v/v) Tween-20 for 1 h at room temperature under gentle agitation. Membranes were washed three times with PBS containing 0.1% (v/v) Tween-20 (PBS-T).

Incubation with primary antibodies was performed overnight (12-14 h) at 4°C using the following antibodies: rat anti-Pt-(G-G/C-G) antibody (ONCOLYZE, Cat. No. R-C18; 1:5,000 dilution), mouse anti-CPD antibody (1:7,500 dilution), and mouse anti-(6-4)PP antibody (1:3,000 dilution), each diluted in PBS-T. After washing three times with PBS-T, membranes were incubated for 1 h at room temperature with HRP-conjugated secondary antibodies diluted 1:5,000 in blocking buffer. An anti-rat IgG-HRP conjugate was used for Pt-(G-G/C-G) detection, whereas an anti-mouse IgG-HRP conjugate was used for CPD and (6-4)PP detection. Chemiluminescent signals were detected using Clarity Western ECL substrate (Bio-Rad, Cat. No. 1705061) and visualized using a Bio-Rad ChemiDoc Imaging System.

To verify equal DNA loading, membranes were stained with SYBR Gold nucleic acid stain (Life Technologies, Cat. No. S11494) diluted 1:5,000 in PBS-T and incubated for 1 h at room temperature in the dark. Membranes were washed three times with PBS-T and imaged using the Bio-Rad imaging system. Signal intensities were quantified using ImageJ software.

#### Histochemical detection of superoxide radicals and hydrogen peroxide

ROS accumulation following UV-B light exposure was evaluated by histochemical staining for superoxide radicals and hydrogen peroxide (H_2_O_2_). For superoxide detection, nitroblue tetrazolium (NBT) staining was performed as previously described [32]. Duckweed plantlets exposed to UV-B light and corresponding controls were collected at 0, 12, and 24 h into recovery. For H_2_O_2_ detection, plantlets were immersed in freshly prepared 0.1% (w/v) 3,3′-diaminobenzidine (DAB) solution adjusted to pH 3.8. Samples were vacuum infiltrated and incubated in the dark at 22°C for 4.5-5 h. Following incubation, chlorophyll was removed by incubating samples in an acetic acid:glycerol:ethanol solution (1:1:3, v/v/v) at 95°C for 10 min [33]. Samples were subsequently stored in 95% (v/v) ethanol until photographed using an Olympus stereomicroscope (SZ61).

## Results

### Effect of UV-B exposure on Spirodela polyrhiza

To evaluate the effects of exposure to UV-B light on *S. polyrhiza*, we treated 13-day-old duckweed plantlets grown *in vitro* with broadband UV-B light (0.2 μmol m^−2^ s^−1^) for 1 min, 2.5 min, 5 min, or 10 min using UVB-313 fluorescent tubes. We assessed morphological responses of untreated control plantlets and those of plantlets following UV-B light treatment. Exposure to UV-B light affected the morphology, coloration, and overall fitness of duckweed plantlets in a duration-dependent manner (Fig. 1). Indeed, plantlets exposed to shorter durations of UV-B light (1 min and 2.5 min) exhibited mild wilting and slight chlorosis relative to the untreated controls. By contrast, plantlets subjected to longer durations of UV-B light (5 min and 10 min) displayed more pronounced damage, characterized by severe wilting, tissue yellowing, and visibly no plant growth. Control plantlets (not exposed to UV-B light) remained healthy-looking and rich in green pigments. These observations suggest that prolonged exposure to UV-B light negatively affects the physiological integrity and pigmentation of *S. polyrhiza*, suggesting greater cellular and photosynthetic damage under extended UV-B stress conditions.

**Fig 1.**
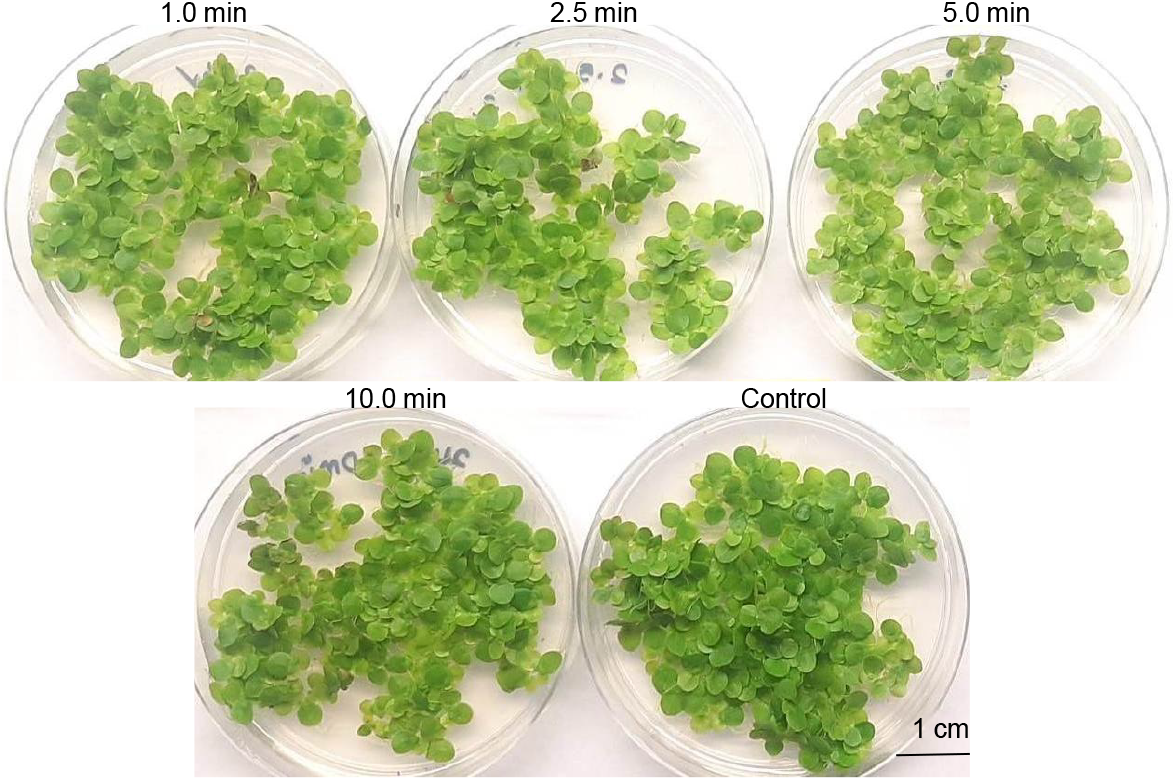
Effect of UV-B exposure on *Spirodela polyrhiza*. Duckweed plantlets were exposed to UV-B radiation (0.2 μmol m^−2^ s^−1^) for 1.0, 2.5, 5.0, and 10.0 min. Samples were photographed immediately after UV-B exposure for 1.0, 2.5, 5.0, and 10.0 min; Control plantlets (not exposed to UV-B light).

### Isolation, quantification, and agarose gel analysis of genomic DNA

To assess the integrity of genomic DNA, we isolated DNA from control *S. polyrhiza* plantlets exposed to UV-B radiation for 1, 2.5, 5, or 10 min, as well as from plantlets allowed to recover under UV-B–free normal growth conditions for 12 or 24 h. Following fluorometric quantification using a Qubit system (Thermo Fisher Scientific), equal volumes consisting of equal amounts of genomic DNA (500 Picograms) from each sample were analyzed by agarose gel electrophoresis. UV-B exposure resulted in a clear, duration-dependent reduction in genomic DNA integrity (Fig. 2). Samples collected immediately after UV-B treatment (0 h recovery) exhibited progressively weaker and less distinct DNA bands compared with the non-irradiated control, with the most pronounced effects observed following 5 and 10 min of exposure. In addition, overall DNA band intensity was markedly reduced in UV-B-treated samples, indicating substantial DNA damage. In contrast, control plantlets not exposed to UV-B displayed strong, intact, high-molecular-weight genomic DNA bands with high signal intensity, consistent with minimal DNA degradation and high DNA quality. Notably, plantlets allowed to recover for 12 or 24 h after UV-B exposure showed a progressive restoration of DNA band intensity relative to the corresponding non-recovered samples. This recovery trend was consistent across all UV-B exposure durations. The gradual increase in DNA integrity during the recovery period suggests activation of DNA repair mechanisms following UV-B-induced damage. Collectively, these results demonstrate that UV-B radiation compromises genomic DNA integrity in *S. polyrhiza* in a dose-dependent manner, while recovery under normal growth conditions promotes partial restoration of DNA integrity, likely through DNA repair processes.

**Fig 2.**
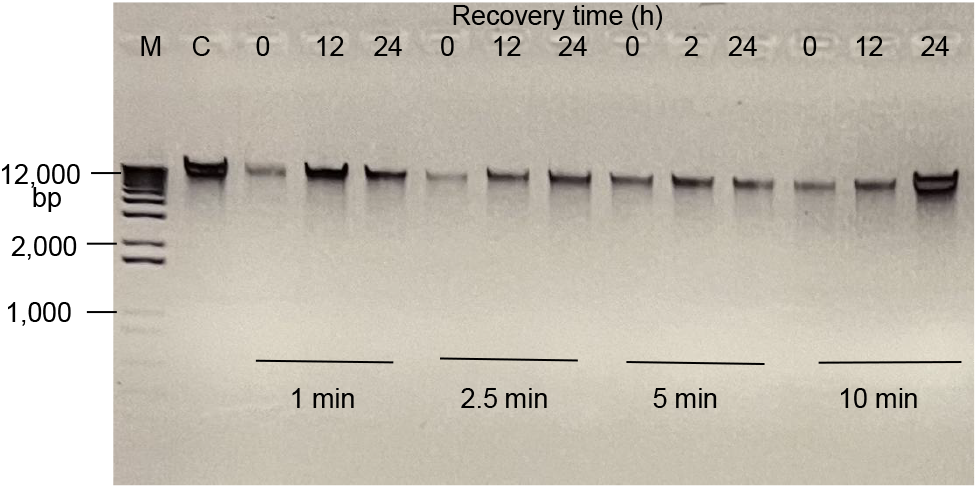
Analysis of UV-B-induced DNA damage and repair in *Spirodela polyrhiza* by agarose gel electrophoresis. Duckweed plantlets were exposed to UV-B radiation (0.2 μmol m^−2^ s^−1^) for 1.0, 2.5, 5.0, and 10.0 min. Samples were collected immediately after UV-B exposure (0 h) and following 12 h and 24 h recovery periods after treatment. Equal amounts of (500 picograms) DNA were loaded in each wells. M, DNA marker; bp, base pair; C, control plantlets (not exposed to UV-B light); 0 h, 12 h, and 24 h indicate recovery periods after UV-B treatment.

### Detection of CPD and 6-4(PP) DNA lesions in UV-B light–exposed duckweed plantlets by immune slot-blot assays

To evaluate UV-B light-induced DNA damage in *S. polyrhiza*, we conducted immuno-slot blot assays using genomic DNA isolated from plantlets exposed to UV-B light for 1 min, 2.5 min, 5 min, or 10 min, alone or after 12 h or 24 h of recovery under UV-B light–free conditions. We used specific antibodies against CPDs (Fig. 3A) or (6-4)PPs (Fig. 3B) to detect UV-B light-induced DNA lesions. The immuno-slot blot analysis revealed the accumulation of both CPD and (6-4)PP lesions in UV-B light–treated plantlets (Fig. 3A and B). Signal intensity increased with longer duration of UV-B light exposure, indicating more pronounced DNA damage under prolonged UV-B stress. By contrast, genomic DNA isolated from untreated control plantlets showed little to no detectable signal for either CPD or (6-4)PP lesions, suggesting minimal basal DNA damage in the absence of UV-B light exposure.

**Fig 3.**
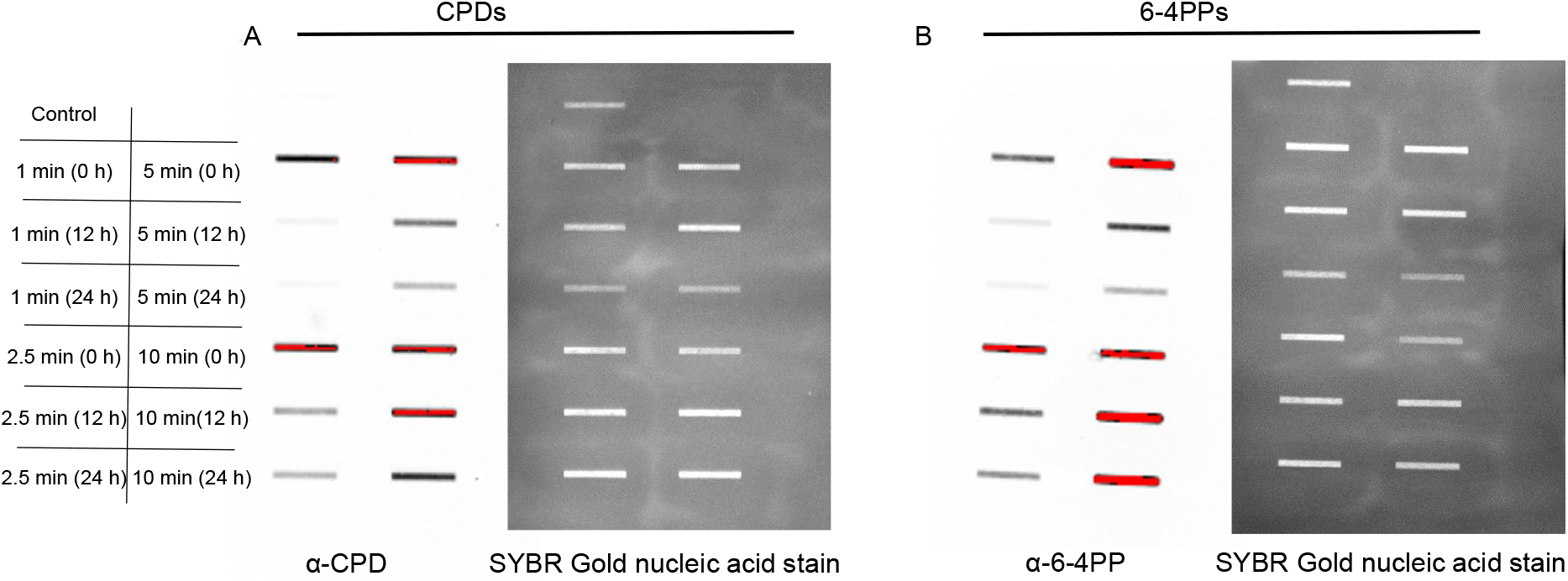
Slot blot analysis of UV-B-induced DNA lesions in *Spirodela polyrhiza*. (A) Detection of cyclobutane pyrimidine dimers (CPDs) after 30 sec. exposure. (B) Detection of 6–4 pyrimidine–pyrimidone photoproducts (6–4PPs) in genomic DNA isolated from duckweed plantlets exposed to UV-B radiation, using an immuno-slot blot assay (30 sec. exposure). Darker bands indicate higher accumulation of CPDs and 6–4PPs, reflecting increased DNA damage. No detectable bands were observed in the control samples, indicating the absence of UV-induced DNA lesions. Duckweed plantlets were exposed to UV-B radiation (0.2 μmol m^−2^ s^−1^) for 1.0, 2.5, 5.0, and 10.0 min. control (plantlets were not exposed to UV-B radiation); 0 h, 12 h, and 24 h indicate recovery periods after UV-B treatment.

To verify equal DNA loading and establish the location of DNA on the membranes, we stained each membrane with SYBR Gold following antibody detection. We measured relative signal intensities (Arbitrary Units, AU) from each slot on the membranes using ImageJ software, and plotted CPD and (6-4)PP levels as a function of UV-B light exposure duration and recovery periods (Fig. 4A and 4B). Samples collected from plantlets exposed to UV-B light but without a recovery period (no recovery) accumulated the highest CPD and (6-4)PP levels. However, lesion signal intensities gradually declined after 12-h and 24-h recovery periods, suggesting active repair of UV-B light–induced DNA damage during post-stress recovery. The decline in CPD and (6-4)PP signals during recovery is consistent with removal of UV-induced photoproducts by endogenous DNA-repair processes; however, because recovery occurred under normal illumination, photoreactivation as well as NER could contribute to lesion removal [12,34,22]. Furthermore, the levels of DNA lesions formed were dependent on the duration of UV-B light exposure, indicating a dose-dependent relationship between UV-B stress and DNA damage responses in S. polyrhiza.

**Fig 4.**
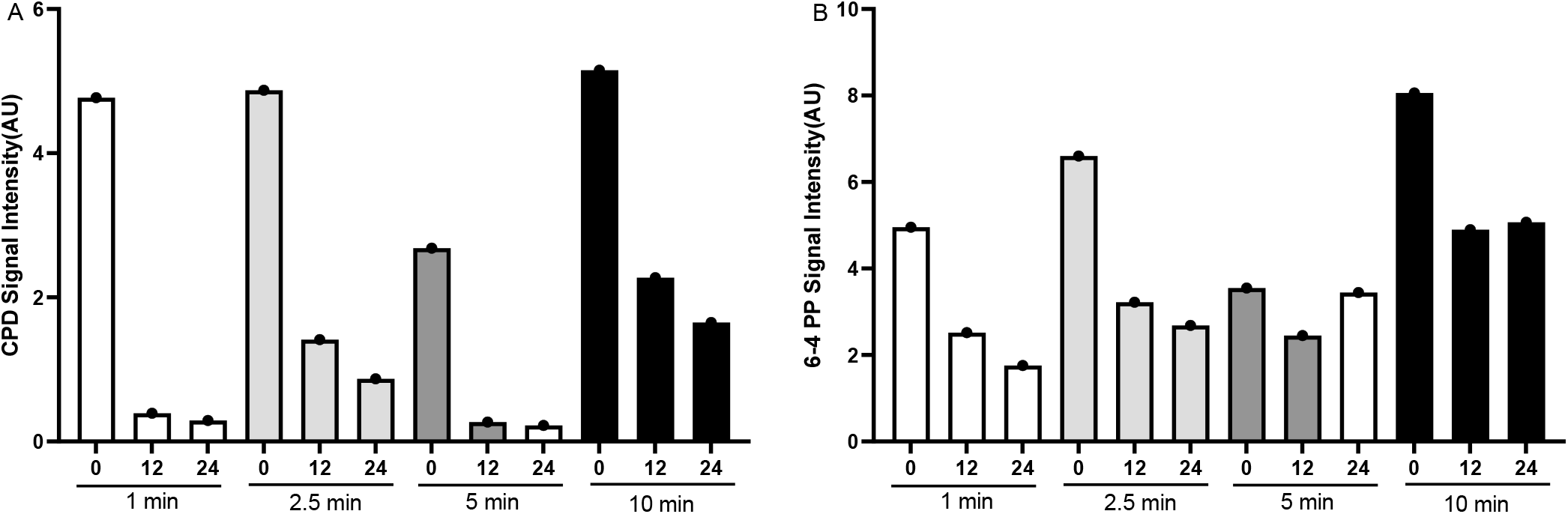
Quantification of UV-B-induced DNA damage and repair in *Spirodela polyrhiza* using ImageJ analysis of immuno-slot blots exposed for 30 sec. (A) Signal intensity of cyclobutane pyrimidine dimers (CPDs). (B) Signal intensity of 6–4 pyrimidine–pyrimidone photoproducts (6–4PPs). Duckweed plantlets were exposed to UVB radiation (0.2 μmol m^−2^ s^−1^) for 1.0, 2.5, 5.0, and 10.0 min. Control plantlets were not exposed to UV-B radiation. Samples were collected immediately after UV-B exposure (0 h) and following 12 h and 24 h recovery periods after treatment (n = 3; average ± SD).

#### In situ visualization of hydrogen peroxide accumulation in UV-B light-exposed duckweed plantlets using DAB staining

We evaluated hydrogen peroxide (H_2_O_2_) accumulation in *S. polyrhiza* plantlets following UV-B exposure by *in situ* 3,3′-diaminobenzidine (DAB) staining. DAB staining revealed substantially higher H_2_O_2_ accumulation in UV-B light-treated plantlets than in untreated controls (Fig. 5). Samples collected from plantlets exposed to UV-B light and allowed no recovery time exhibited intense dark-brown staining, indicating accumulation of this ROS following UV-B stress. By contrast, control plantlets (not exposed to UV-B light) showed minimal staining, suggesting low basal H_2_O_2_ levels under normal growth conditions. The amount of DAB staining rose with longer durations of exposure to UV-B light, suggesting a dose-dependent oxidative stress response. Plantlets exposed to 5 min or 10 min of UV-B light displayed strong staining intensity and thus high H_2_O_2_ accumulation within their fronds. Notably, we observed a lower staining intensity in samples collected after 12 h or 24 h of recovery than in the corresponding no-recovery samples. This diminished H_2_O_2_ accumulation during the recovery phase suggests activation of antioxidant defense and stress recovery mechanisms following UV-B light exposure. Collectively, these findings indicate that UV-B light induces oxidative stress in *S. polyrhiza* by promoting H_2_O_2_ accumulation in a dose-dependent manner, whereas post-treatment recovery facilitates partial alleviation of oxidative damage over time.

**Fig 5.**
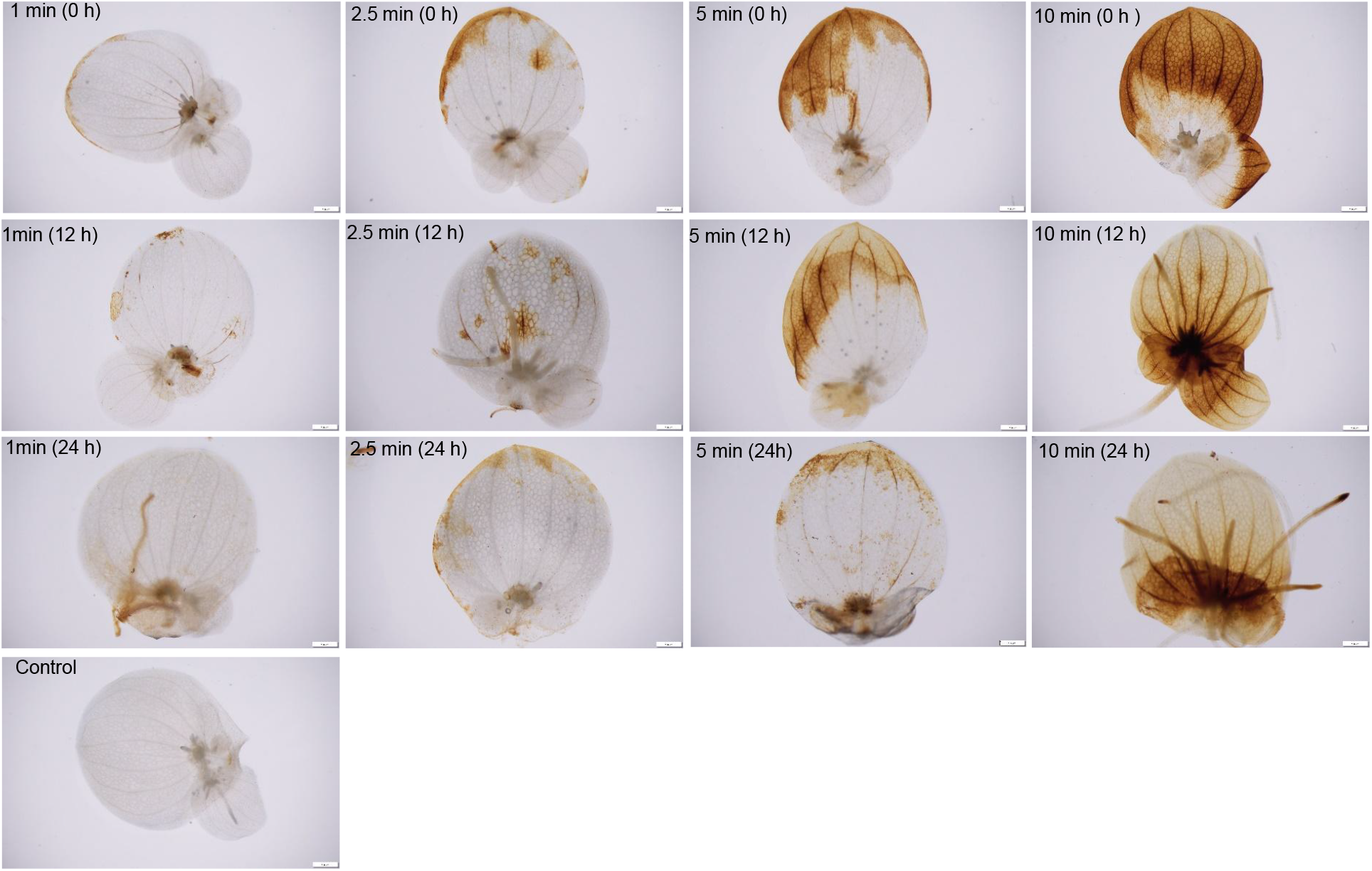
In situ visualization of hydrogen peroxide (H_2_O_2_) accumulation in *Spirodela polyrhiza* using DAB (3,3-diaminobenzidine) staining. DAB staining revealed a higher accumulation of H_2_O_2_ in duckweed plantlets immediately after UV-B exposure (0 h recovery period), as indicated by intense brown coloration. In UV-B-treated plantlets, H_2_O_2_ accumulation progressively decreased during the 12 h and 24 h recovery periods. No detectable H_2_O_2_ accumulation was observed in the control plantlets that were not exposed to UV-B radiation.

## Discussion

In the present study, we demonstrate that UV-B light induces substantial DNA damage and oxidative stress in the aquatic monocotyledonous plant *S. polyrhiza*, while simultaneously activating DNA repair-associated regulatory pathways during post-stress recovery. The observed accumulation of CPDs, (6-4)PPs, and H_2_O_2_, collectively indicate that duckweed possesses an active and coordinated defense system to mitigate UV-B light-induced stress. These findings support the utility of *S. polyrhiza* as a promising experimental model for investigating plant DNA damage and repair mechanisms under UV-B stress.

Morphological analysis revealed that prolonged exposure to UV-B light gradually impaired *S. polyrhiza* growth, pigment accumulation, and overall fitness. Plantlets exposed to longer periods of UV-B light exhibited chlorosis, tissue wilting, and lower fitness, all common physiological symptoms associated with UV light-induced oxidative and photosynthetic damage in plants. Similar UV-B-associated growth inhibition, pigment changes, and photosynthetic impairment have been documented across plant systems, reflecting the combined effects of photodamage, oxidative stress, and acclimation responses [8,11,22]. Because *S. polyrhiza* fronds possess minimal tissue complexity and are directly exposed to surrounding environmental conditions, they may be particularly sensitive to UV-B light-mediated oxidative injury. The dose-dependent morphological changes observed in this study therefore likely reflect cumulative cellular damage associated with longer (10 min) exposure to UV-B light.

Consistent with these phenotypic observations, genomic DNA integrity was markedly affected by UV-B light. Agarose gel electrophoresis showed lower intensity for large DNA bands in UVB light-treated samples, particularly following longer exposure to UV-B light, suggesting greater DNA fragmentation or structural disruption. Importantly, DNA integrity appeared to return to some extent following recovery under UV-B light-free conditions for 12 h or 24 h, suggesting the activation of endogenous DNA repair pathways. These findings are consistent with recent work showing that plants deploy photoreactivation, NER, BER, translesion synthesis, and double-strand-break repair to remove UV-associated lesions and preserve genome stability [23,24,22].

We detected direct evidence of UV-B light-induced DNA damage in the form of the accumulation of CPD and (6-4)PP lesions by immuno-slot blot assays. Both lesion types accumulated rapidly following exposure to UV-B light and in a clear dose-dependent manner. CPDs and (6-4)PPs are the major UV-induced DNA photoproducts formed after absorption of UV photons by pyrimidine-rich DNA; these lesions can impede replication and transcription [16,18,22]. These lesions interfere with DNA replication and transcription by distorting the DNA helical structure and blocking the progression of DNA polymerase. Similar UV-B light–induced accumulation of CPDs and (6-4)PPs has been extensively documented in mammals and plants, including Arabidopsis, maize (*Zea mays*), and rice [9,35,36]. Notably, lesion abundance declined during recovery, particularly after 24 h, indicating active repair of UV light-induced DNA damage in *S. polyrhiza* tissues.

The progressive decline in CPD and (6-4)PP levels during recovery strongly suggests the involvement of the NER and photoreactivation pathways in *S. polyrhiza*. In plants, CPDs are primarily repaired through photolyase-mediated photoreactivation under light conditions, whereas (6-4)PP lesions are repaired by both photolyases and NER-associated mechanisms [20,12]. Because recovery in this study occurred under UV-B light-free normal illumination provided by a growth chamber, visible light-dependent photolyase activity likely contributed substantially to lesion removal. In addition, the gradual drop in DNA lesion levels over time suggests that *S. polyrhiza* possesses efficient UV-responsive DNA repair systems comparable to those previously characterized in land plants [37,34,38].

UV-B light exposure also triggered substantial oxidative stress in *S. polyrhiza* plantlets, as evidenced by the high accumulation of H_2_O_2_ detected by DAB staining. We observed the strongest H_2_O_2_ accumulation in samples that were not allowed to recover under normal light conditions following prolonged UV-B light exposure. ROS are important mediators of UV-B stress and can contribute to oxidative damage of proteins, lipids, nucleic acids, and membranes while also functioning as signals that regulate stress acclimation [10,11,22]. Elevated ROS production under UV-B stress has previously been linked to impaired electron transport, chloroplast dysfunction, and disruption of cellular redox homeostasis in plants [39]. The decline in DAB staining intensity during recovery is consistent with the activation of antioxidant defense systems that mitigate oxidative stress after UV-B light exposure. Although we did not directly measure the activity of antioxidant enzymes in the present study, previous studies have demonstrated that UV-B stress is associated with higher bulk activity for antioxidant enzymes such as superoxide dismutase, catalase, and ascorbate peroxidase, which detoxify ROS and restore cellular homeostasis [9,10].

The present findings demonstrate that UV-B light induces both direct DNA photo damage and secondary oxidative stress in *S. polyrhiza*, while simultaneously activating repair and recovery mechanisms that contribute to restoration of cellular integrity (Fig. 6). The relatively rapid decline seen in the levels of DNA lesions during recovery highlights the efficiency of duckweed DNA repair systems under controlled environmental conditions. Moreover, the small genome size, rapid vegetative propagation, simple morphology, and ease of laboratory cultivation make *S. polyrhiza* an attractive model system for studying UV light-induced DNA damage responses. Future studies integrating transcriptomics, proteomics, antioxidant enzyme analyses, and functional characterization of additional DNA repair genes should improve our understanding of UV-B light tolerance mechanisms in duckweed and may provide broader insights into plant acclimation to ozone layer damage and UV light associated with global environmental change.

**Fig 6.**
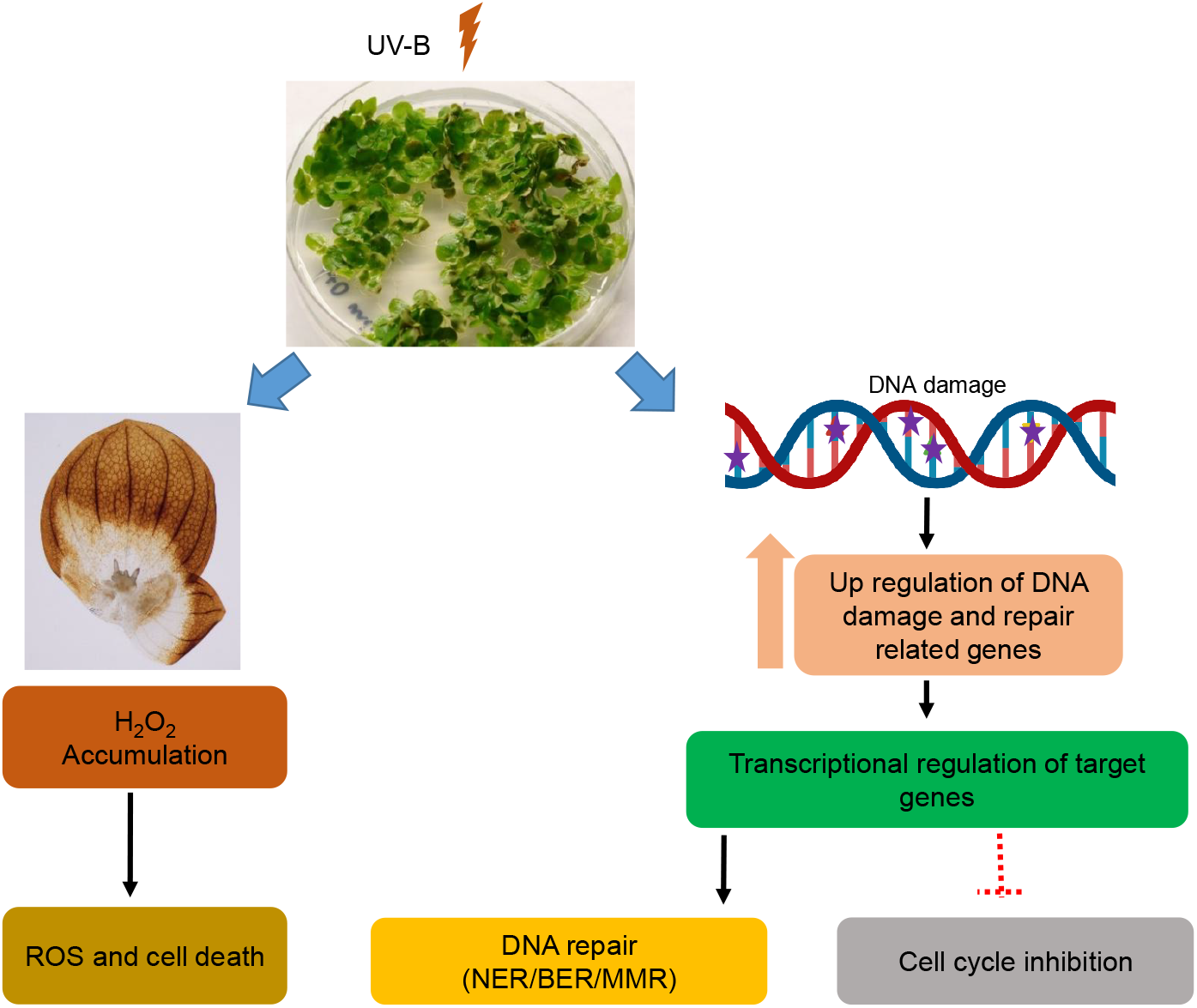
Schematic overview of the effects of UVB light stress in *Spirodela polyrhiza*. The schematic illustrates the induction of reactive oxygen species (ROS) and UV-B-induced DNA damage in duckweed, leading to the activation of DNA damage response and repair pathways.

## Acknowledgements

We thank Professor. Shobhan Gaddameedhi and Dr. Venugopal Bovilla, North Carolina State University, for generously providing the CPD and (6-4)PP antibodies and for their expert guidance and mentorship through virtual discussions and video meetings.

